# Fluctuations in populations of subsurface methane oxidizers in coordination with changes in electron acceptor availability

**DOI:** 10.1101/040204

**Authors:** C. Magnabosco, P.H.A. Timmers, M.C.Y. Lau, G. Borgonie, B. Linage-Alvarez, O. Kuloyo, R. Alleva, T.L. Kieft, G.F. Slater, E. van Heerden, B. Sherwood Lollar, T.C. Onstott

## Abstract

The concentrations of electron donors and acceptors in the terrestrial subsurface biosphere fluctuate due to migration and mixing of subsurface fluids, but the mechanisms and rates at which microbial communities respond to these changes are largely unknown. Subsurface microbial communities exhibit long cellular turnover times and are often considered relatively static—generating just enough ATP for cellular maintenance. Here, we investigated how subsurface populations of CH_4_ oxidizers respond to changes in electron acceptor availability by monitoring the biological and geochemical composition in a 1,339 meters-below-land-surface (mbls) fluid-filled fracture over the course of both longer (2.5 year) and shorter (2-week) time scales. Using a combination of metagenomic, metatranscriptomic, and metaproteomic analyses, we observe that the CH_4_ oxidizers within the subsurface microbial community change in coordination with electron acceptor availability over time. We then validate these findings through a series of ^13^C-CH_4_ laboratory incubation experiments, highlighting a connection between composition of subsurface CH_4_ oxidizing communities and electron acceptor availability.

## Introduction

The terrestrial subsurface is an energy-limited environment that is subject to changes in fluid chemistry over time (Onstott *et al.* 2006). Laboratory experiments have shown that when native fluids are supplemented with electron acceptors such as SO_4_^2-^, the activity of subsurface communities can be enhanced (Rajala *et al.* 2015). Large disturbances such as CO_2_ (Morozova *et al.* 2010, 2011) and H_2_ injection (Bagnoud *et al.* 2016), hydraulic fracking (Daly *et al.* 2016), and drilling (Purkamo *et al.* 2013) have also been reported to alter natural subsurface communities. The response of microbial communities to natural fluctuations in their environment, however, is less understood.

In the South African subsurface, increases in the availability of electron acceptors such as NO_3_^−^ (>10-fold higher) and SO_4_^2-^ (>5-fold higher) over a 1.5 year period did not correspond to changes in the bacterial community (Magnabosco *et al.* 2014; Simkus *et al.* 2015). However, 16S SSU rRNA gene amplicon surveys of the archaeal communities from the same site and sampling points of the aforementioned studies showed a diverse collection of anaerobic methane oxidizing archaea (ANME) (Young *et al.* 2017) and methane oxidizing bacteria (Simkus *et al.* 2015).

ANME-1 and “*Candidatus* Methanoperedens nitroreducens”, a member of ANME-2d, are among the best described ANME and couple the oxidation of CH_4_ with SO_4_^2-^ and NO_3_^−^ reduction, respectively (Haroon *et al.* 2013; Wegener *et al.* 2015). Other ANME-2d have also been reported to couple CH_4_ oxidation to the reduction of Mn^4^+ and/or Fe^3^+ (Ettwig *et al.* 2016). With the exception of “*Ca*. Methylomirabilis oxyfera” which has been suggested to generate intracellular O2 for CH_4_ oxidation from two molecules of NO (Ettwig *et al.* 2010), bacterial methanotrophs couple CH_4_ oxidation with free O2 in the environment. This potential relationship between CH_4_ oxidizers (MOs) and electron acceptor availability provides a compelling avenue to explore the response of subsurface communities to natural changes in subsurface fluid chemistry.

Our study focuses on the subsurface microbial community of a 1,339 meters-below-land-surface (mbls) fluid-filled fracture (Be326 Bh2). Here, bulk bacterial phospholipid-derived fatty acid (PLFA) isotopic signatures have been shown to be consistent with the accumulation of ^13^C-dissolved inorganic carbon (DIC) impacted by the microbial oxidation of CH_4_ (Simkus *et al.* 2015) but the organisms responsible for CH_4_ oxidation have not been well characterized (Magnabosco *et al.* 2014; Simkus *et al.* 2015; Young *et al.* 2017). To better describe the membership of native MO communities, their methane oxidizing genes, and their response to changes in fluid chemistry over time, we combined metagenomics, metatranscriptomics, and metaproteomics analyses with geochemical monitoring of Be326 Bh2’s *in situ* fracture fluids over both 2.5 year and 2 week timescales. We also performed activity assays on the fracture fluids using ^13^C-labeled CH_4_ together with different electron acceptors to track anaerobic methane oxidizing activity in both short- and long-term incubations.

## Materials and Methods

Be326 Bh2 was accessed through a 57-m, horizontally drilled borehole that was drilled in 2007 and sealed with a high-pressure steel valve. The borehole is located on the 26 level of shaft 3 in the Beatrix Gold Mine (28.232288 S, 26.794365 E; Welkom, South Africa). Annual samples were collected during field trips in 2011, 2012, and 2013 and weekly samples were collected during the 2013 field trip with three time points designated as T_0_, T_1_, and T_2_ with T_0_ corresponding to the first day of the 2013 study and T_2_ corresponding to the last day of the 2013 study.

### Sampling

Sampling procedures have been outlined previously (Magnabosco *et al.* 2014). To sample, a sterile (combusted and autoclaved) stainless steel manifold with attached valves was attached to the Be326 borehole. A high-pressure steel valve was opened, allowing water to flow freely at ~4-6 L min^−1^ for at least 10 minutes. This manifold allows for the attachment of airtight Teflon tubing and filters for sampling and chemical analysis. For the 2011, 2012, and 2013 samples, pre-autoclaved 0.1-μm Memtrex NY filters (MNY-91-1-AAS; General Electric Co., Minnetonka, MN USA) were left on the borehole for 6, 15, and 14 days, respectively. For the T_0_, T_1_, and T_2_ samples, 0.2-μm Memtrex NY Capsule (CMNY) filters (General Electric Co., Minnetonka, MN USA) were left on the borehole for 2 hours with a flow rate of 500 mL per minute (equivalent to approximately 60 L filtered per time point) in 2013. The total volumes of water filtered for each time point in the 2.5-year study were 4,875 L, 12,604 L, and 6,635 L in 2011, 2012 and 2013, respectively.

Unfiltered water samples for direct cell counts were collected on the first day of the 2011, 2012, and 2013 studies and fixed with sterile formaldehyde (final concentration 4% v/v). Cell counts were obtained at the University of the Free State where samples were filtered through a sterile 0.22μm Millipore GTTP-type membrane filter, stained with DAPI, and visualized using fluorescence microscopy.

### Geochemical Measurements

Temperature, pH, and *Eh* were measured at the borehole using handheld probes (HANNA instruments, Woonsocket, RI). Gas samples (H2, O2, CH_4_) were collected and analyzed by gas chromatography (Peak Performer 1 series, Peak Laboratories, USA) (Simkus *et al.* 2015). Oxygen was also measured at the borehole using a CHEMET kit (Chemetrics Inc.; Calverton, VA). NO_3_^−^ and SO_4_^2-^ were measured using an ion chromatograph coupled to an electrospray ionization-quadruple mass spectrometer (MS) (Dionex IC25 and Thermo Scientific MSQ, USA). δ^18^O and δ^2^H were measured as previously described (Simkus *et al.* 2015).

### Preservation of Biomolecules and extraction

Filters were treated with an RNA preserve solution and stored in a −80°C freezer. The RNA preserve is a custom made solution of 20 mM EDTA, 0.3 M sodium citrate, and 4.3 M ammonium SO_4_^2-^ (pH 5.2). The solution was autoclaved prior to sample application. Total protein, together with total DNA and RNA, was extracted using 2X CTAB lysis buffer and phenol/chloroform (pH=6.5-6.9), and re-suspended in 1 TE-buffer (Tris-EDTA, pH = 8) and stored in 1.5-mL Eppendorf tubes at −20°C. Extraction of biomolecules is further described elsewhere (Lau *et al.* 2014, 2016).

### Amplicon Sequencing and Annotation

The V6 region of archaeal 16S rDNA molecules from the 2011 and 2012 time points was amplified using 958F (AATTGGANTCAACGCCGG) and 1048R (CGRCRGCCATGYACCWC) primers. The V4-V5 region of archaeal 16S rDNA molecules from all time points was amplified using the 517F (GCCTAAAGCATCCGTAGC; GCCTAAARCGTYCGTAGC; GTCTAAAGGGTCYGTAGC; GCTTAAAGNGTYCGTAGC; GTCTAAARCGYYCGTAGC) and 958R (CCGGCGTTGANTCCAATT) primers. For both amplicon datasets, forward primers included 5-nt multiplexing barcodes and a reverse 6-nt index. Triplicate PCR amplifications were performed in 33-μL reaction volumes with an amplification cocktail containing 10.0 U Platinum Taq Hi-Fidelity Polymerase (Invitrogen, Carlsbad, CA), 1X Hi-Fidelity buffer, 200 μM dNTP PurePeak DNA polymerase mix (ThermoFisher), 2 mM MgSO_4_ and 0.3 μM of each primer. We added approximately 10-25 ng template DNA to each PCR and ran a control without template DNA for each primer pair. Amplification conditions were: initial 94°C, 3 minute denaturation step; 25 cycles of 94°C for 30 s, 60°C for 45 s, and 72°C for 60 s; and a final 2 minute extension at 72°C. The triplicate PCR reactions were pooled after amplification and purified using Qiagen MinElute plates followed by clean up, PicoGreen quantitation and Sage PippinPrep size selection. 101-nt paired-end sequencing was performed on an Illumina HiSeq 1000 at the Marine Biological Laboratory (Woods Hole, MA USA). Only reads that were identical in the overlapping regions of the forward and reverse reads were included for annotation. In the case of V6 data, this filter necessitates an exact match across both the forward and reverse read. Sequences that met the given quality criteria were annotated using GAST (Huse *et al.* 2008) and a GAST-formatted reference set.

### Mapping of Metatranscriptomic Data to V6 Amplicons

Quality filtered RNA reads were mapped to the GAST-annotated V6 amplicons of Be326 Bh2 (Magnabosco *et al.* 2014; Young *et al.* 2017) using BLASTn (percent identity >97%; alignment length > 55 nucleotides (nt)). Reads that mapped positively to V6 sequences were assigned a consensus taxonomy based on the top three BLASTn hits.

### Generation of Metagenomic and Metatranscriptomic Datasets

DNA samples from 2011 and 2012 were sequenced at the National Center for Genome Resources (Santa Fe, NM). The 2011 and 2012 metagenomic libraries were prepared from 500 ng of DNA using the KAPA High Throughput Library Preparation Kit (KAPA Biosystems), an insert size of approximately 280 bp, and followed by 8 PCR cycles. Paired-end sequencing (2 × 100 nt) was performed on an Illumina HiSeq 2000. DNA from 2013 was sequenced at Lewis Sigler Institute for Integrative Genomics (Princeton, NJ USA) using an Illumina HiSeq 2500. The metagenomic library was prepared using the TruSeq Rapid SBS Kit, size selected for 380-400 nt, and sequenced using paired-end sequencing (2 × 215 nt).

RNA samples from 2011, 2012, 2013, T_0_, T_1_, and T_2_ were sequenced at Lewis Sigler Institute for Integrative Genomics (Princeton, NJ USA) on an Illumina HiSeq 2500 platform. Here, metatranscriptomic libraries were prepared using the Ovation RNA-Seq v2 System (NuGEN; San Carlos, CA USA), which involved 15-18 cycles of PCR. The 2011 and 2012 RNA samples were sequenced using 141-nt single-end sequencing while the 2013, T_0_, T_1_, and T_2_ RNA samples were sequenced using 200-nt single-end reads.

### Targeted Assembly and Tree Building

Reads were quality filtered and assembled using a targeted assembly pipeline (https://github.com/cmagnabosco/OmicPipelines). In summary, this pipeline involved quality filtering reads using default settings on the fastx toolkit (http://hannonlab.cshl.edu/fastx_toolkit/), targeted assembly of *mcrA* GENEs, protein prediction, alignment of reads to a set of well curated McrA peptides, and the construction of a phylogenic tree from the predicted proteins (PhyML (Guindon *et al.* 2010), Best of nearest-neighbor-interchange and subtree-pruning-regrafting, 8 rate categories, −b 100). This procedure was also repeated for the assembly of *mmo* genes using a MMO peptide database.

Assembled genes of 200 nt or longer were trimmed to include only the coding region of verified *mcrA* and *mmo* genes. This dataset was used as the reference database (Supplementary Data 2) to map quality-filtered DNA and RNA reads using Bowtie2 (--very-sensitive-local). The coverage was calculated as: (number of reads mapped × average read length)/length of the reference sequence. For correlation analyses, coverage was normalized by dividing by the total number of reads in the dataset. Pearson correlation coefficients (r) were computed in Excel using the CORREL function.

### Protein Identification from UPLC-MS/MS Data

Ultra performance liquid chromatography-tandem mass spectrometer (UPLC-MS/MS) data were analyzed in aggregate using the SEQUEST HT search engine in ProteomeDiscoverer v1.4 (ThermoFisher Scientific) using a custom database containing the translated genes from the targeted assembly and published McrA, pMoA, MmoX and AmoA (Supplementary Data 4). Search parameters included trypsin digestion with up to one missed cleavage, methionine oxidation and cysteine carbamidomethylation. A peptide-level false discovery rate (FDR) of 5% was achieved by using the Percolator node in ProteomeDiscoverer, which utilizes the frequency of matching against a reversed database as a rigorous model of the probability of error in the forward matches at given score thresholds. Proteins identified by matches to one unique peptide per run were tallied.

### Activity assays

During the 2012 and 2013 sampling points, water samples were collected into 150-mL serum vials (“biovials”) and stored at 4°C. Two days prior to sampling, 100 μL of MilliQ water was added to a 150-mL serum vial that had undergone combustion in an oven at 450°C overnight to deactivate spores. The serum vial was then sealed with a butyl rubber stopper that was cleaned by boiling in 0.1 M NaOH solution for 1 hour, rinsed and left to soak in MilliQ water until use. The serum vials were then crimped and left overnight. The next day, the serum vials were purged and pressurized with filtered N2. The vials were then autoclaved and bubble-wrapped for transport underground. A needle was attached to the end of sterile teflon tubing that was attached to the manifold. Water was allowed to flow to rinse the attachment and needle. After 2 minutes of rinsing, the needle was inserted into the biovial and a second needle was added for pressure relief. Water was allowed to flow into the biovial until the relief needle overflowed. Extensive care was taken to ensure that no visible gas bubble remained in the biovial post-sampling.

Two sets of enrichment experiments were performed to monitor the response of the biovial communities to methane and a variety of electron acceptors. Experiment 1 contained samples from 2012 and 2013. Samples were incubated in triplicate with ^13^C-CH_4_ and SO_4_^2-^ (20 mM) or ^13^C-CH_4_ and NO_3_^−^ (20 mM) for 207 and 185 days, respectively. A control incubation with ^13^C-CH_4_ and no electron acceptors was monitored for 207 days to measure endogenous ^13^CO_2_ production (Control A). Experiment 2 included samples from 2012 and 2013 that were incubated in triplicate with ^13^C-CH_4_ and SO_4_^2-^ (20 mM), NO_3_^−^ (20 mM) or O2 (5% v/v). A killed (paraformaldehyde, 4% v/v) control without electron acceptors (Control 1) and a live control without electron acceptor or donor were included (Control 2). All samples of Experiment 2 were incubated and monitored over a 43-day period.

The samples for all experiments were prepared in 14-mL serum vials sealed with butyl rubber stoppers and aluminum caps. Prior to sample addition, vials were made anoxic by exchanging headspace gas with N2 for 10 cycles and left pressurized (0.5 bar overpressure). 10 mL aliquots of fracture fluid were then added to the vials in an anoxic chamber and amended with the treatments described above. When ^13^C-CH_4_ was added, N_2_ was added to a pressure of 130 kPa and 99.99% ^13^C-CH_4_ gas (Campro Scientific, Veenendaal, The Netherlands) was added to a final pressure of 180 kPa. Oxygen was added afterwards, when applicable. All electron acceptor solutions were sterile and anoxic. The serum bottles were statically incubated at 37°C in the dark.

NO_3_^−^ and SO_4_^2-^ were analysed using an ion chromatography system equipped with an Ionpac AS9-SC column and an ED 40 electrochemical detector (Dionex, Sunnyvale, CA). The system was operated at a column temperature of 35°C, and a flow rate of 1.2 ml min^−1^. Eluent consisted of a carbonate/bicarbonate solution (1.8 and 1.7 mM respectively) in deionized water. Headspace gas composition was measured on a gas chromatograph-mass spectrometer (GC-MS) composed of a Trace GC Ultra (Thermo Fisher Scientific, Waltham, MA) equipped with a Rt-QPLOT column (Restek, Bellefonte, PA), and a DSQ MS (Thermo Fisher Scientific). Helium was used as a carrier gas with a flow rate of 120 ml min^−1^ and a split ratio of 60. The inlet temperature was 80°C; the column temperature was set at 40°C and the ion source temperature was 200°C. CH_4_ and CO_2_ in the headspace were quantified from the peak areas in the gas chromatographs. The fractions of ^13^CO_2_, ^13^CH_4_ and ^12^CH_4_ were derived from the mass spectrum (Shigematsu *et al.* 2004). Validation of the method was done using standards with known mixture of ^13^CO_2_ and ^12^CO_2_. The concentrations of total CO_2_, total CH_4_, and ^13^CO_2_ were calculated following the method of Timmers *et al.* (2015). The pressure of the serum vials was determined using a portable membrane pressure unit (GMH 3150, Greisinger electronic GmbH, Regenstauf, Germany). The pH was checked by a standard pH electrode (QiS, Oosterhout, The Netherlands).

## Results and Discussion

### A changing fluid chemistry over time

Over a period of 2.5 years, water isotope analysis revealed large changes in the fracture fluid’s δ^2^H and δ^18^O isotopic signatures (Fig. 1; purple squares). These changes are inconsistent with contamination with service water in the mine and, instead, indicate mixing of different fracture waters within the system. In 2011, the δ^18^O and δ^2^H values were on the global meteoric water line (GMWL) and are indicative of paleometeoric water. This isotopic signature is consistent with other fluids located approximately 1,000 to 1,500 mbls in the Witwatersrand Formation (South Africa) (Fig. 1; green triangles) (Onstott *et al.* 2006; Sherwood Lollar *et al.* 2007). In 2012 and 2013, the δ^18^O and δ^2^H signatures of the fluids moved away from the GMWL, indicating mixing with more ancient fluids (Frape *et al.* 1984). A few meters away, Be326 Bh1 (a borehole located in the same mine and depth as the Be326 borehole of this study) exhibited a similar trend in δ^18^O and δ^2^H signatures over time—shifting away from the GMWL in 2012 and 2013 (Fig. 1; red diamonds). The displacements between the 2011/2012 and 2012/2013 δ^18^O and δ^2^H signatures of the two boreholes are in opposite directions—a pattern that would not be expected if the native fracture fluids were mixing with the mine’s service water during this period. Measurements of the fracture fluid’s δ^2^H and δ^18^O isotopic signatures over the 2-week study were not obtained.

**Figure 1.**
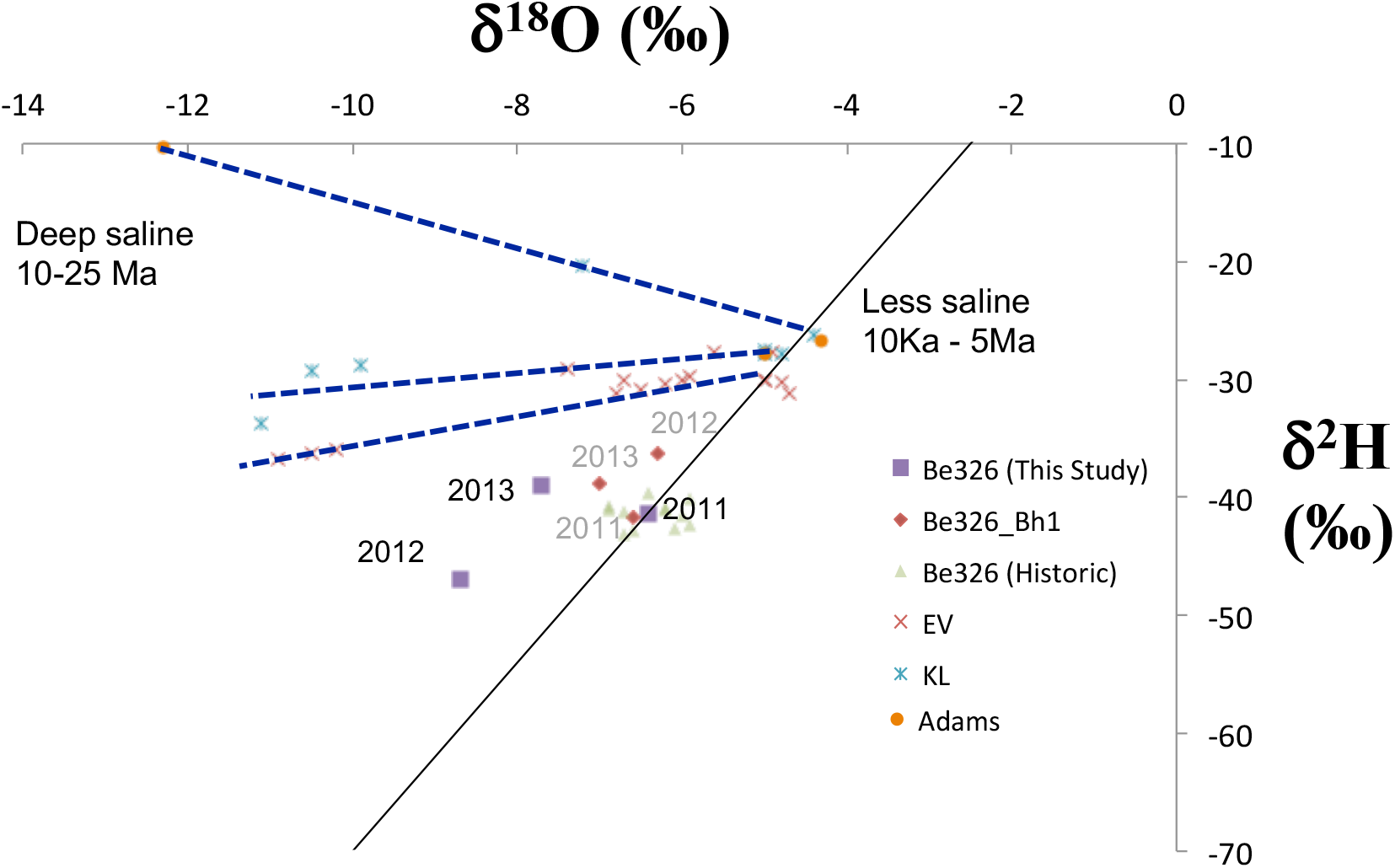
δH_H_2_O_ and δ^18^O_H_2_O_ for Beatrix (Be) Gold Mine fracture fluids. This figure plots the δ^18^0 (x-axis) against the δ^2^H (y-axis) for each of the three time points (purple squares) in the long-term study of Be326 (2011, 2012, 2013). The black diagonal line represents the global meteoric water line (Craig 1961) (GMWL). The 2011 sample sits on the GMWL, consistent with historic Be datapoints (green triangles) and paleometeoric water (Lippmann *et al.* 2003; Ward *et al.* 2004). The 2012 and 2013 time points are displaced away from the GMWL along trends that have been observed to be consistent with mixing with older, more saline fracture fluids (Ward *et al.* 2004; Sherwood Lollar *et al.* 2007). The dotted blue lines indicate possible mixing patterns. Be326_Bhl (red diamonds) is a borehole located in the same mine and at the same depth as the borehole sampled in this study. EV=Evander Mine; KL=Kloof. Data for EV, KL, and Adams are adapted from Ward *et al.* (2004).

Geophysico-chemical measurements were made for both the 2.5-year and 2-week time series (Table 1). Temperature, pH, and CH_4_ concentrations were relatively unchanged in both of the time series but the degree to which E_h_, SO_4_^2-^, NO_3_^−^, and H_2_ concentrations changed was much greater over the 2.5 years (Table 1a). Within the 2.5-year time series, the fracture fluids shifted from a more reduced state (2011_E*h*_ = −98 mV, 2011_[H2]_ = 130 nM) with limited electron acceptor availability (2011_[Sulph.]_ = 137μM, 2011_[Nitr.]_ = 0.4 μM) to a more oxidized state (2013_E*h*_ = 21±28 mV, 2013_[H2]_ = 25 nM) with much greater electron acceptor availability (2013_[Sulph.]_ = 479 μM, 2013_[Nitr.]_ = 4.5 μM). The 2-week time series did not exhibit as large of a shift in electron acceptor availability as the 2.5-year time series and maintained a positive E_*h*_ throughout (Table 1b).

**TABLE 1a.** Geophysico-chemical measurements for the long-term study

|  | **2011** | **2012** | **2013** |
| --- | --- | --- | --- |
| **Dates Sampled** | Jan 21-27, 2011 | July 12-27, 2012 | Aug 1-15, 2013 |
| **Temperature (°C)** | 36.9 | 37.3±1.2 | 35.1±0.8 |
| **pH** | 8.83 | 8.55 | 8.17±0.7 |
| **E<sub>h</sub> (mV)** | -98 | -27.5±6.4 | 21±28 |
| **SO<sub>4</sub><sup>2-</sup> (μM)** | 137 | 623 | 479 |
| **NO<sub>3</sub><sup>-</sup> (μM)** | 0.4 | 6.0 | 4.5 |
| **O<sub>2</sub> (Chemet) (μM)** | b.d. | 0.47 | b.d.* |
| **H<sub>2</sub> (nM)** | 130 | 9 | 25 |
| **CH<sub>4</sub> (mM)** | 2.0 | 0.9 | 1.1 |

b.d. = below detection (<0.31 μM)
b.d.\* = below detection (<1.6 μM)

**TABLE 1b.** Geophysico-chemical measurements for the short-term study

|  | **T<sub>0</sub>** | **T<sub>1</sub>** | **T<sub>2</sub>** |
| --- | --- | --- | --- |
| **Dates Sampled** | Aug 1, 2013 | Aug 8, 2013 | Aug 15, 2013 |
| **Temperature (°C)** | 35.8 | 35.3 | 28.6 |
| **pH** | 8.2 | 8.1 | 7.9 |
| **E<sub>h</sub> (mV)** | 9 | 53 | 1 |
| **SO<sub>4</sub><sup>2-</sup> (μM)** | 531 | 499 | 459 |
| **NO<sub>3</sub><sup>-</sup> (μM)** | 5.3 | 5.3 | 3.7 |
| **O<sub>2</sub> (Chemet) (μM)** | b.d.* | b.d.* | b.d.* |
| **H<sub>2</sub> (nM)** | 5.8×10<sup>-5</sup> | 10.4 | 39.8 |
| **CH<sub>4</sub> (mM)** | 0.8 | 1.2 | 1.0 |

b.d.\* = below detection (<1.6 μM)

### Microbial community of Be326

The microbial communities of the 2011 and 2012 time points have been reported to be dominated by bacteria (98.5%) (Simkus *et al.* 2015) with the majority being related to Proteobacteria (Magnabosco *et al.* 2014). In order to investigate the community composition of the less numerous archaea, 16S rDNA amplicon sequencing of the archaeal V4-V5 hypervariable region was performed across all time points (Fig. 2a). For the long-term study (2011, 2012, 2013), the archaeal community shifted from an ANME-1- and Methanomicrobia-dominated community in 2011 to a Miscellaneous Crenarchaeotic Group-dominated community in 2012 and a Halobacteria-dominated community in 2013. For the short-term study (T_0_, T_1_, T_2_), there were no noticeable changes between T_1_ and T_2_. A slight increase in Halobacteria and a decrease in ANME-1 with respect to T_1_ and T_2_ were observed in the T_0_ time point. There is a noticeable difference between the T_0_, T_1_, T_2_ samples community profiles and the 2013 community profile, despite being collected over the same two-week period. However, the filters used in the Tx and 2013 filtrations have varied pore sizes, geometries, and casings and should not be directly compared.

**Figure 2.**
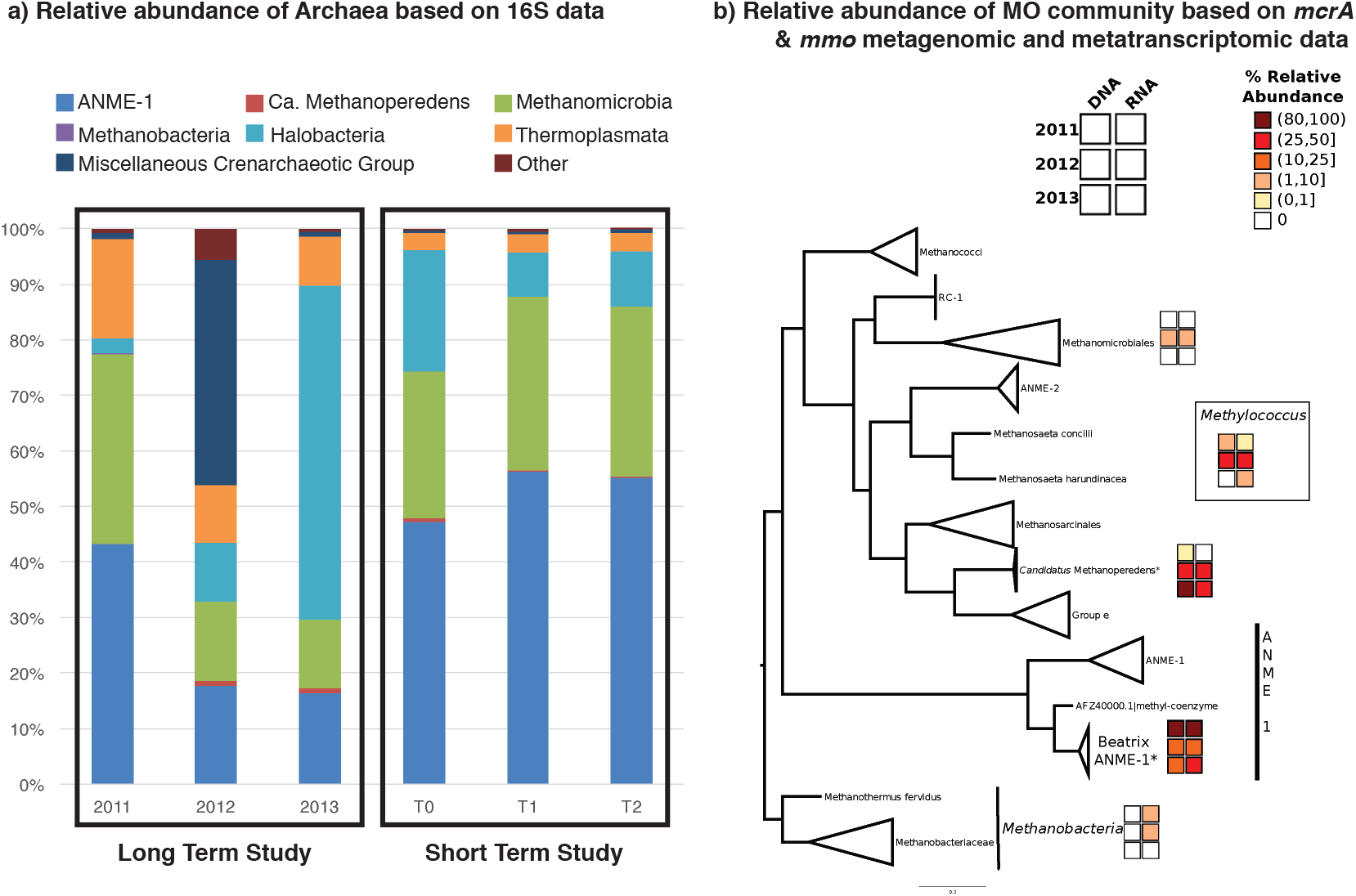
Change in the CH_4_ oxidizing community and activity over time. a) The relative abundance (bar height) of major archaeal groups (colors of bars) identified in the archaeal V4-V5 16S rDNA surveys are shown. b) A phylogenetic tree was constructed using PhyML from a McrA peptide database (Supplementary Data 6) and the predicted proteins from the *mcrA* targeted assembly (designated by a *). The predicted MMO from the targeted assembly is also displayed. The 2×3 blocks represent the relative abundance of each taxon (DNA, left column; RNA, right column) with respect to their CH_4_-related protein of interest (McrA or MMO) over time (row 1, 2011; row 2, 2012; row 3; 2013). Notably, Methanobacteria-related McrA PEGs were selected as the outgroup in this presentation of the McrA phylogenetic tree due to Methanobacteria’s placement in species tree of Euryarchaeota.

Around 1% of V4-V5 amplicons were related to “*Ca.* M. nitroreducens” in all time points except for the 2011 dataset that contained only 2 archaeal amplicons relating to *“Ca.* M. nitroreducens” (Supplementary Data 1). These relative abundances of “Ca. M. nitroreducens” are lower than what was reported using archaeal V6 primers on 2011 and 2012 samples. With archaeal V6 primers, *“Ca.* M. nitroreducens” accounts for 1.5% of the archaeal community in 2011 and 10.6% of the archaeal community in 2012, while ANME-1 accounts for ~10% of the archaeal community at each time point (Young *et al.,* 2017; Supplementary Fig. S1). Despite differences in the relative abundances of taxa based on the primers used, community membership does not appear to be significantly different between the 2 primer sets.

In order to estimate the relative activity of each taxonomic group, metatranscriptomic data from each sample were mapped to a database of Be326 Bh2 16S rDNA V6 bacterial (Magnabosco *et al.* 2014) and archaeal (Young *et al.* 2017) sequences. The V6 sequences were selected over the V4-V5 sequences due to their shorter length and stringent quality filtering procedure (see Materials & Methods). Proteobacteria within the family Rhodocyclaceae dominated all of the RNA datasets except for the 2011 dataset that was dominated by Hydrogenophilaceae. The number of bacterial and archaeal species (unique hits within the 16S rDNA V6 database) observed in the metatransciptomic data ranged from 204 in 2013’s T0 time point to 414 in the 2012 sample (Table 2). MO archaea and bacteria account for only a small percentage of the V6 rRNA sequences identified in the metatranscriptomic data (0.3-3%). Notably, the number of species observed in each metatranscriptome’s V6 rRNA pool was not correlated to the number of reads generated for each metatranscriptome (R^2^ < 0.01); however, the number of species observed in each metatranscriptome is almost 10 times less than the number of OTU0.97 obtained from the bacterial 16S rDNA V6 dataset (2,478 in 2011 and 3,987 in 2012) (Magnabosco *et al.* 2014).

**Table 2:** Metatranscriptomic species richness observed and estimated by Chao 1 and ACE metrics and cell concentrations measured in the field.

|  | 2011 | 2012 | 2013 | T<sub>0</sub> | T<sub>1</sub> | T<sub>2</sub> |
| --- | --- | --- | --- | --- | --- | --- |
| **Species (S) Observed** | 302 | 414 | 247 | 222 | 204 | 248 |
| **S.Chao1** | 344±22 | 440±10 | 256±7 | 233±9 | 216±8 | 276±16 |
| **S.ACE** | 317±8 | 450±10 | 253±6 | 233±5 | 212±7 | 260±8 |
| **Cell concentration (×10<sup>3</sup> cells mL<sup>-1</sup>)** | 1.3<sup>^</sup>-<br>18.3<sup>*</sup> | 2.2<sup>*</sup>-<br>31<sup>^</sup> | 3.2<sup>^</sup> | 3.2<sup>^</sup> | n.m. | n.m. |

<sup>^</sup>DAPI-derived cell concentration
<sup>\*</sup>PLFA-derived cell concentration using a conversion factor of 6×10<sup>4</sup> cells/pmol PLFA as previously reported in Simkus et al. (2015)
n.m. = not measured

### A detectable shift in the MO community over time

As the organisms responsible for CH_4_ oxidation in Be326 were present at relatively low abundances, a targeted assembly pipeline (https://github.com/cmagnabosco/OmicPipelines) that employs the PRICE assembler (Ruby, Bellare and DeRisi 2013) was implemented to assemble methyl-coenzyme M reductase *(mcrA*)—the gene for the first step in the anaerobic oxidation of methane (AOM) (Thauer 2011) or the last step in methanogenesis—and a suite of CH_4_ monooxygenases *(mmo)* that are known to play a role in aerobic oxidation of methane (McDonald *et al.* 2008) from the metagenomic and metratranscriptomic datasets. Notably, *mcrA* was selected as an indicator for ANME presence because its phylogeny is congruent with MO phylogeny.

Following targeted assembly and annotation, two complete *mcrA* genes related to ANME-1 and “Ca. M. nitroreducens”, an ANME-2d, were assembled. Only one complete *mmo* gene closely related to *Methylococcus capsulatus* was recovered from the metagenomic and metatranscriptomic data (Fig. 2b, Supplementary Data 2). Partial *mcrA* related to Methanomicrobia and Methanobacteria were also identified in the high-throughput data (Fig. 2b, Supplementary Data 2), but partially assembled mmo-related genes were omitted in downstream analyses due to the difficulty in distinguishing *mmo* from homologous ammonia monooxygenases genes (Holmes *et al.* 1995).

Assembled *mcrA* and *mmo* were translated into peptide sequences (Supplementary Data 3). Notably, the Be326 *“Ca.* M. nitroreducens”-type McrA was 99% identical to the McrA of the reference *“Ca.* M. nitroreducens” genome (Supplementary Data 3, 4). Metaproteomic data were searched against the collection of assembled McrA and MMO and a database of known McrA and MMO peptide sequences (Supplementary Data 4) to confirm that the transcribed genes were translated into proteins. The predicted amino acid sequences (Supplementary Data 3) from the assembled ANME-1, *“Ca.* M. nitroreducens”, and *Methylococcus* genes were all identified within the metaproteomic data (Fig. 3, Supplementary Data 5) which further confirmed the presence and activity of these groups of organisms.

**Figure 3:**
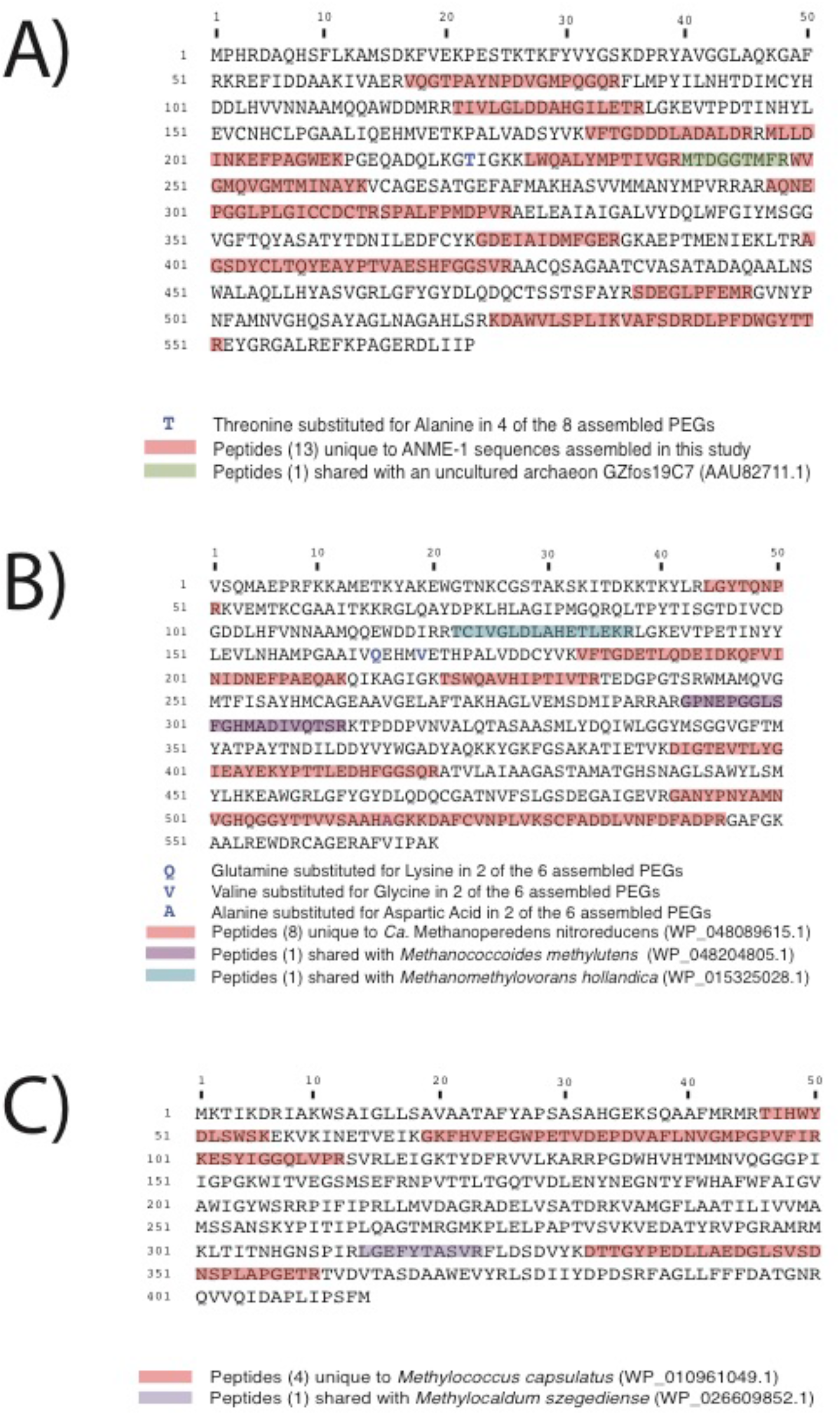
Peptide mapping to identified McrA and MMO proteins.

The abundances (based on coverage) of each MO gene *(mcr,A mmo)* and MO 16S SSU rRNA gene were calculated using Bowtie2 (Langmead and Salzberg 2012) (-very-sensitive-local) and BLASTn, respectively. These analyses provided evidence that the dominant members of the MO community in the metagenomes and metatranscriptomes shifted from ANME-1 to *“Ca.* M. nitroreducens” during the 2.5-year sampling period (Fig. 2) but remained constant during the 2-week sampling period (Supplementary Table 1). These observations were consistent with the relative changes in geochemistry over both time scales (Table 1). Notably, *“Ca.* M. nitroreducens” accounted for ≤1% of the archaeal community, as revealed using archaeal V4-V5 16S rDNA primers (Fig. 1), which is in contrast to the higher estimates of *“Ca.* M. nitroreducens” found when using archaeal V6 16S rDNA (Young *et al.,* 2017; Supplementary Fig. S1), metagenomic and metatranscriptomic data. This discrepancy suggests that *“Ca.* M. nitroreducens” sequences are recovered at a lower efficiency in archaeal V4-V5 16S SSU rRNA gene surveys relative to archaeal V6 16S SSU rRNA gene surveys, metagenomic and metatranscriptomic studies.

When metagenomic MO *mcrA* and *mmo* abundances of the long-term study (Supplementary Table 2) were correlated to geophysico-chemical measurements, *“Ca.* M. nitroreducens” was positively correlated to NO_3_^−^ (R^2^=0.99) and SO_4_^2-^ (R^2^=0.98) concentrations but negatively correlated to CH_4_ (R^2^=0.99) and H_2_ (R^2^=0.99) concentrations. ANME-1 *mcrA* abundances showed an opposite trend and were positively correlated to CH_4_ (R^2^=0.96) and H_2_ (R^2^=0.97) concentrations but negatively correlated NO_3_^−^ (R^2^=0.88) and SO_4_^2-^ (R^2^=0.85) concentrations (Table 3). Correlation of metatranscriptomic 16S rRNA and *mcrA* and *mmo* abundances to geophysico-chemical measurements exhibited similar trends (Supplementary Table 3) and are consistent with a transition from an ANME-1-dominated MO community to a *“Ca.* M. nitroreducens”-dominated community. As NO_3_^−^-coupled CH_4_ oxidation is more energetic than SO_4_^2-^-coupled CH_4_ oxidation (Caldwell *et al.* 2008), an energetic advantage, presumably, provides *“Ca.* M. nitroreducens” a competitive advantage against ANME-1 when NO_3_^−^ concentrations are sufficient.

**Table 3:** Correlation (R^2^) between normalized metagenomic *mcrA* or *mmo* PEG coverage and geochemical parameters. Positive correlations are underlined.

|  |  | NO<sub>3</sub><sup>-</sup> | SO<sub>4</sub><sup>2-</sup> | CH<sub>4</sub> | *E<sub>h</sub>* | pH | H<sub>2</sub> |
| --- | --- | --- | --- | --- | --- | --- | --- |
| **2.5-Year**<br>**Time Series** | “*Ca. Methanoperedens*” | <u>0.99</u> | <u>0.98</u> | 0.99 | <u>0.70</u> | 0.52 | 0.99 |
|  | ANME-1 | 0.88 | 0.85 | <u>0.93</u> | 0.90 | <u>0.76</u> | <u>0.95</u> |
|  | *Methylococcales* | <u>0.12</u> | <u>0.14</u> | 0.07 | 0.10 | <u>0.25</u> | 0.05 |

O_2_ concentrations in 2011 and 2013 were below detection limit (Table 1) but were detectable in 2012 (0.47 μM). Likely related to the increased availability of O2, aerobic *Methylococcus-related mmo* genes exhibited their highest relative abundances within metagenomic and metatranscriptomic MO gene profiles during 2012 and a minimal presence throughout the remainder of the time points (Fig. 2, Supplementary Table 1). “Ca. M. nitroreducens” was the dominant member (73.2±2.8%) of the MOs community during the 2-week time series (Supplementary Table 1) when fracture fluids contained high concentrations of SO_4_^2-^ (496±36 μM) and NO_3_^−^ (4.8±0.9 μM) along with a positive Eh (21+28 mV).

The correlations between MO abundances and fluid chemistry suggest that a relationship between electron acceptor availability and populations of MOs exists. We therefore wanted to experimentally validate the response of the MOs to changes in electron acceptor availability. As ANME-1, “Ca. M. nitroreducens”, and *Methylococcus* are best described as a SO_4_^2-^-dependent ANME (Wegener *et al.* 2015), NO_3_^−^-dependent ANME (Haroon *et al.* 2013), and aerobic methanotrophs (Kleiveland *et al.* 2012), respectively, we designed experiments to test whether or not each MO lifestyle would respond to an increase in the aforementioned electron acceptor.

### Validation of the MO community through ^13^CH_4_ enrichments

To better understand the response of the MO community to changes in electron donor/acceptor balance, two sets of ^13^C-CH_4_ laboratory enrichment experiments were performed on fracture water collected in 2012 and 2013. The first experiment (Experiment 1) was a long-term experiment analyzed over 207^*^ days and contained fracture fluid samples from 2012 and 2013 enriched with either ^13^C-CH_4_ and no additional electron acceptor (endogenous activity control), ^13^C-CH_4_+SO_4_^2-^ (to stimulate SO_4_^2-^-dependent AOM), or ^13^C-CH_4_+NO_3_^−^ (to stimulate NO_3_^−^-dependent AOM). A second set of 2012 and 2013 fracture fluid enrichments (Experiment 2) was analyzed for 43 days. Experiment 2 contained ^13^C-CH_4_ treatments of ^13^C-CH_4_+formaldehyde (4%, v/v) (killed control) to rule out non-biological sources of ^13^C-CH_4_ production, ^13^C-CH_4_+SO_4_^2-^, ^13^C-CH_4_+O2 (to simulate aerobic methane oxidation), and ^13^C-CH_4_+NO_3_^−^ as well as an electron acceptor- and donor-free control (methanogenesis control). The methanogenesis control was intended to detect whether or not methanogenesis, and therefore also trace CH_4_ oxidation (TMO),was occurring within the samples (Zehnder and Brock 1979). Due to the limited amount of sample, we were unable to test the potential occurrence of Mn^4^+-or Fe^3^+-driven methane oxidation.

Unlike the correlations observed between expression data and geochemical parameters, an increase in the proportion of ^13^C-CO_2_ relative to total CO_2_ (%^13^C-CO_2_) in the ^13^CH_4_ enrichments over time provides definitive evidence of ^13^CH_4_ oxidation under different conditions. Notably, we chose to express our results as %^13^C-CO_2_ rather than the absolute concentration (molar) of ^13^C-CO_2_ to account for CO_2_ production from other substrates. In Experiment 1, the ^13^C-CH_4_+NO_3_^−^ enrichments exhibited the greatest rate of %^13^C-CO_2_ production and, in 2012, the rate of %^13^C-CO_2_ production (0.017±0.005 %^13^C-CO_2_ day^−1^) was found to be significantly greater (paired one-tailed Student t-test; p=0.02) than in Control A (0.004±0.001 %^13^C-CO_2_ day^−1^) (Fig. 4). No samples from Experiment 2 exhibited an increase in %^13^C-CO_2_ production (Supplementary Data 5).

**Figure 4:**
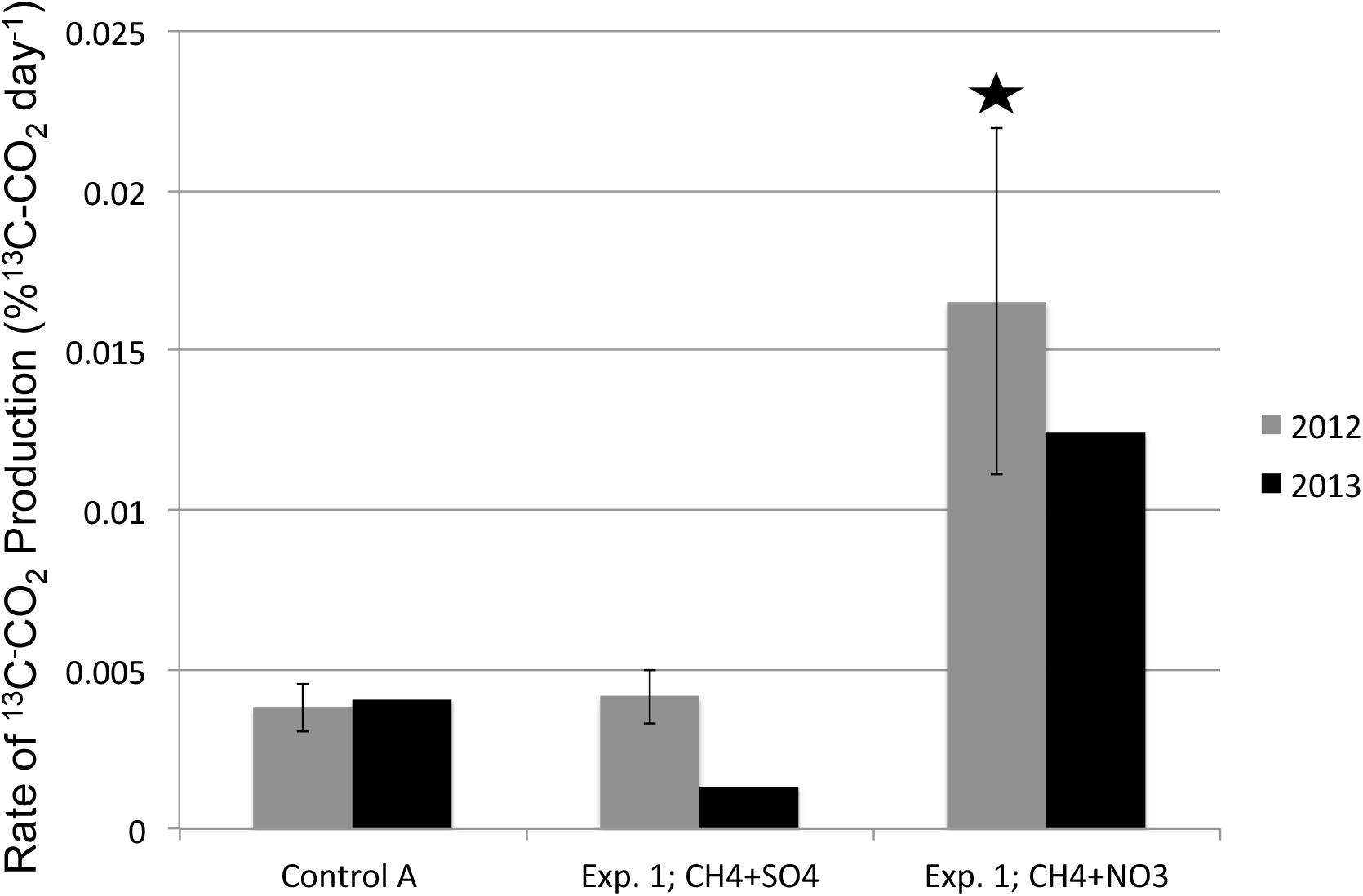
^13^C-CO_2_ production after ^13^C-CH_4_ enrichment. The rates of ^13^C-CO_2_ production (%^13^C-CO_2_ day^−1^) from the 2012 (gray) and 2013 (black) fluids of Experiment 1 are displayed. The 2012 samples were run in triplicate and the standard deviations are shown. The star above the 2012 CH_4_+NO_3_^−^ bar indicates that the rate of ^13^C-CO_2_ production in the ^13^C-CH_4_+NO_3_^−^ enrichment was significantly greater (*p*=0.02) than the electron acceptor-free control (Control A).

Although ANME-1 were present in the metatranscriptomic and metaproteomic data during the 2012 and 2013 sampling points, there was not a significant difference in %^13^C-CO_2_ production between Experiment 1’s ^13^C-CH_4_+SO_4_^2-^ incubations and the endogenous activity controls (Fig. 4). TMO was probably not responsible for the ^13^C-CO_2_ production in the endogenous activity controls of Experiment 1; only the methanogenesis control of the 2013 sample showed minor methanogenesis (and thus TMO) activity (Supplementary Data 5). It is conceivable, however, that SO_4_^2-^-dependent AOM occurred in both the endogenous controls and ^13^C-CH_4_+SO_4_^2-^ incubations, as the concentrations of SO_4_^2-^ in the controls ([SO_4_^2-^ *2012*] = 623 μM; [SO_4_^2-^ *2013*] = 479 μM) are well within the lower range of SO_4_^2-^ concentrations (100-1200 μM) that have been reported for SO_4_^2-^-coupled AOM (Beal, Claire and House 2011; Segarra *et al.* 2015). Combined, these findings suggest that SO_4_^2-^-coupled AOM likely occurred in the controls of Experiment 1 and 2.

## Conclusions

Metagenomic, metatranscriptomic, and metaproteomic data suggest that community composition, activity, and function are changing in response to natural fluctuations in fluid chemistry. The observed CH_4_ oxidation in the controls and dominance of ANME-1 in the 2011 samples (when SO_4_^2-^ concentrations were lowest) indicate that the *in situ* fluids contain enough SO_4_^2-^ to power SO_4_^2-^-coupled MO. The increase in %^13^C-CO_2_ production in the ^13^C-CH_4_+NO_3_^−^ enrichments and correlation of *“Ca.* Methanoperedens” abundances to electron acceptor concentrations *in situ* suggest that electron acceptor availability plays an important role in MO population dynamics. Together, these results provide the most conclusive biological evidence to date that CH_4_ oxidation occurs and is an integral component of the deep terrestrial subsurface carbon cycle.

## Data Availability

Metagenomic and metatranscriptomic data are available at NCBI BioProject PRJNA308990. 16S amplicon data are available under NCBI BioProject PRJNA263371.

## Supplement

Supplementary Figure S1 – Relative abundance of archaeal 16S SSU rRNA gene v6 amplicons for the 2011 and 2012 time points

Supplementary Table 1 – CH_4_ oxidizing community with respect to their C¾-related gene

Supplementary Table 2 – Normalized coverage C¾-related genes abundances Supplementary Table 3 – Correlation of CH_4_ oxidizing community activity to fluid chemistry

Supplementary Data 1 – Archaeal V4-V5 16S rDNA annotations

Supplementary Data 2 – Assembled *mcrA* and *mmo* genes

Supplementary Data 3 – Predicted proteins from assembled *mcrA* and *mmo*

Supplementary Data 4 – Datasets used for protein mapping

Supplementary Data 5 – Protein mapping results

Supplementary Data 6 – Enrichment experiment results

Supplementary Data 7 – McrA protein library used to generate phylogenetic tree

## Acknowledgments

This research was supported by funding from the National Science Foundation to T.C.O. (EAR-0948659) and to TLK (EAR-0948335) and the Deep Carbon Observatory (Alfred P. Sloan Foundation) to M.C.Y.L. (Sloan 2013-10-03, subaward 48045). Research of P.H.A.T. is supported by the Soehngen Institute of Anaerobic Microbiology (SIAM) Gravitation grant (024.002.002) of the Netherlands Ministry of Education, Culture and Science and the Netherlands Organisation for Scientific Research (NWO). Partial support for isotopic analyses was provided by the Natural Sciences and Engineering Research Council of Canada. We are indebted to the logistical support of Sibanye Gold Limited, the management and staff of Beatrix Gold Mine and specifically to S. Maphanga of Beatrix gold mine. We thank Matthew Cahn (Department of Molecular Biology, Princeton University) and the staff of Research Computing (Office of Information Technology, Princeton University), especially Robert Knight, for their technical support with the computational analyses.

## Footnotes

* The long term ^13^C-CH_4_+NO_3_” enrichments were run for 183 days.

